# Tubulin tyrosination/detyrosination regulates the sorting of intraflagellar transport trains on axonemal microtubule doublets

**DOI:** 10.1101/2024.05.03.592312

**Authors:** Aditya Chhatre, Ludek Stepanek, Adrian Pascal Nievergelt, Gonzalo Alvarez Viar, Stefan Diez, Gaia Pigino

## Abstract

Assembly and function of cilia rely on the continuous transport of ciliary components between the cell body and the ciliary tip. This is performed by specialized molecular machines, known as Intraflagellar Transport (IFT) trains. Anterograde IFT trains are powered by kinesin-2 motors and move along the B-tubules (enriched in detyrosinated tubulin) of the ciliary microtubule doublets. Conversely, retrograde IFT trains are moved by dynein-1b motors along the A-tubules (enriched in tyrosinated tubulin) back to the cell body. The segregation of oppositely directed trains on A-tubules or B-tubules is thought to prevent traffic jams in the cilium, but the mechanism by which opposite polarity trains are sorted onto either tubule, and whether tubulin tyrosination/detyrosination plays a role in that process, is unknown. Here, we show that CRISPR-mediated knock-out of VashL, the enzyme that detyrosinates microtubules, causes recurrent stoppages of IFT trains and reduces the rate of ciliary growth in *Chlamydomonas reinhardtii*. To test whether the observed stoppages, potentially caused by collisions between oppositely directed IFT trains, are ascribable to direct interactions between IFT trains and tubulin tyrosination/detyrosination, we developed methods to reconstitute the motility of native IFT trains from cilia on de-membranated axonemes and *ex vivo* on synthetically polymerized microtubules. We show that anterograde trains have higher affinity for detyrosinated microtubules (analogous to B-tubules), while retrograde trains for tyrosinated microtubules (analogous to A-tubules). We conclude that tubulin tyrosination/detyrosination is required for the spatial segregation of oppositely directed trains and for their smooth uninterrupted motion. Our results provide a model for how the tubulin code governs molecular transport in cilia.

## Introduction

Cilia (or eukaryotic flagella) are conserved microtubule-based organelles of eukaryotic cells that play fundamental roles in fluid motility, cell swimming, and signalling. For their assembly and maintenance they require specialized molecular bi-directional transport machines, the intraflagellar transport (IFT) trains [1]. IFT trains consist of large polymeric assemblies of two large protein complexes known as IFT-A (6 proteins) and IFT-B (16 proteins) [2]. They are powered by microtubule-based molecular motors. Specifically, kinesin-2 and dynein-1b (dynein-2 in humans), respectively, drive the anterograde (base-to-tip) and retrograde (tip-to-base) IFT trains. Thereby, the motion of each IFT train is restricted to a very narrow space just underneath the ciliary membrane along the external face of the nine microtubule doublets that compose the ciliary axonemal structure [1]. Despite the restricted and crowded environment, collisions between oppositely directed IFT trains and consequent traffic jams are not observed in motile cilia. This is because opposite polarity trains sort onto distinct tubules of the microtubule doublets [3], effectively clearing the way for oncoming trains. Retrograde trains move on the A-tubule, while anterograde trains on the B-tubule of the doublet [3]. However, the mechanism responsible for the sorting of the IFT trains onto the different tubules remains elusive.

Though being composed of different numbers of protofilaments, A-tubules and B-tubules have the same tubulin-lattice structure[4]. They are made of the same tubulin isoforms, but have been shown to be differentially enriched in certain tubulin post-translational modifications [5]. Thus, it has been speculated that tubulin post-translational modifications might encode different biochemical cues on the surface of the two tubules that contribute to the spatial segregation of anterograde and retrograde IFT trains. Tubulin polyglutamylation was shown to be enriched in B-tubules [5] and we have recently shown that this post-translational modification specifically marks only a single tubulin protofilament (B9) [6]. Polyglutamylation contributes to the regulation of the Nexin-Dynein Regulatory Complex (N-DRC) function in keeping the axoneme together during ciliary beating [6], [7]. However, protofilament B9 is in an area of the microtubule that is inaccessible for IFT trains, thus unlikely to contribute to IFT regulation. Accordingly, depletion of glutamylation did not affect IFT motility [7]. Tubulin glycylation was instead found on subsets of protofilaments on both A-tubule and B-tubule that can be used by IFT trains [6]. Thus, while glycylation has a role in regulating axonemal dynein activity [8], and might affect IFT motors in general, it is unlikely to contribute to the mechanism of selection of either A-tubule or B-tubule by the trains. Tubulin detyrosination, on the other hand, was shown to be strongly enriched on the B-tubule of Chlamydomonas cilia by immunogold-EM [9] and sequential purification of tubulin from microtubule doublets [5], but its role in cilia remains elusive.

In mammals, it was recently discovered that the genetically encoded C-terminal tyrosine of alpha-tubulin is cleaved post-translationally after microtubule polymerization by tubulin carboxypeptidases such as the Vasohibin-SVBP complex [10] or MATCAP [11]. The Chlamydomonas homolog of these enzymes is not annotated, but detyrosination of axonemal microtubules is a conserved feature across ciliated species. *In vitro* studies have shown that mammalian kinesin-1 [12], [13], [14] and kinesin-2 [15] exhibit higher landing affinity and processivity on detyrosinated microtubules. Additionally, cytoplasmic dynein complexes have preferential affinity for tyrosinated microtubules via the CAP-Gly domain of its adaptor dynactin microtubule binding domain [16], [17], [18]. Thus, the observed differential distribution of tubulin tyrosination and detyrosination could encode signals on the A-tubule and B-tubule that might be read by IFT motors and contribute to the spatial segregation of retrograde and anterograde IFT trains, respectively.

In this study, we show that the absence of tubulin detyrosination in the cilia of Chlamydomonas mutants resulted in recurrent stoppages of IFT trains. Concurrent with these stoppages and alteration of IFT dynamics, we observed a reduction in the ciliary growth rate. The absence of structural defects in the mutant cilia, as revealed by cryo-electron tomography (cryo-ET), suggested a direct interaction between the IFT trains and tubulin detyrosination. To test this hypothesis, we developed a method to reconstitute the motility of native IFT trains on de-membranated cilia (parent axonemes) and *ex vivo* on synthetically polymerized microtubules enriched for different post-translational modifications. We found that anterograde and retrograde IFT trains have differential affinities for detyrosinated and tyrosinated tubulin. Based on our results we conclude that the enrichment of detyrosinated tubulin on the B-tubule [5], [9] biases the sorting of IFT trains, by selectively recruiting anterograde trains while excluding retrograde trains. In the mutant cells, the absence of tubulin detyrosination removes this bias and induces the missorting of the IFT trains onto the wrong microtubules. This gives rise to disruptive train collisions that critically influence IFT dynamics and seamless transport of cargo within cilia.

## Results

### Depletion of tubulin detyrosination in Chlamydomonas VashL mutant cells leads to IFT train stoppages and slows down ciliary growth

To test whether tubulin detyrosination plays a role in regulating IFT train motility in Chlamydomonas, we used CRISPR [19] to generate a mutant with impaired tubulin detyrosination activity. We identified Cre05.g241751_4532 as a suitable locus to disrupt in the Chlamydomonas genome, as it aligns to the SVBP (NCBI RefSeq: NP_955374.1) peptide of the human Vasohibin-SVBP complex (see Materials and Methods) (**Fig. S1**). This insertional CRISPR knock-out mutant (further referred to as *VashL*) exhibited a strong reduction in the levels of detyrosinated tubulin (**Fig. S2A**). We performed the knock-out on wild-type cells as well as on a background strain containing fluorescently tagged IFT46 to study the dynamics of IFT trains in the absence of tubulin detyrosination by TIRF microscopy.

IFT trains in the *VashL* cells showed altered dynamics compared to wild-type cells. Trains exhibited a recurrent ‘stop-and-go’ movement, that is, fast runs were interrupted by transient stationary phases (**Fig. 1A**, green or magenta arrowheads, see also **Fig. S3**). These stoppage events predominantly correlated with the crossing of opposite-polarity trains (**Fig. 1A**, white arrowheads), suggesting that in the *VashL* cells anterograde and retrograde trains experience collisions. Quantification individual stoppage events and normalization relative to total crossing events of each train revealed that anterograde trains were more likely to stop than retrograde trains after a crossing (**Fig. 1B**). This effect was independent of the total number of trains per cilium, as the train injection rates were similar in *VashL* and wild-type cells (**Fig. 1C**). Interestingly, anterograde trains stopped 8% more often in *VashL* cells than in wild-type cells (**Fig. 1B**). This value matches the theoretical probability of collisions for opposite-polarity IFT trains randomly-distributed over all available tubules; the chance that a retrograde train physically interacts with a given anterograde train at a crossing relies on being on the same of the two tubules of the same doublet (1/2 x 1/9 = 0.5 x 11.11% ≅ 6%). While stoppages due to collisions are unlikely in wild-type cells (as opposite polarity trains do not walk on the same tubule of any microtubule doublet [2]), the observed increase in the stoppage events in the *VashL* cells thus points towards the presence of collisions.

**Figure 1.**
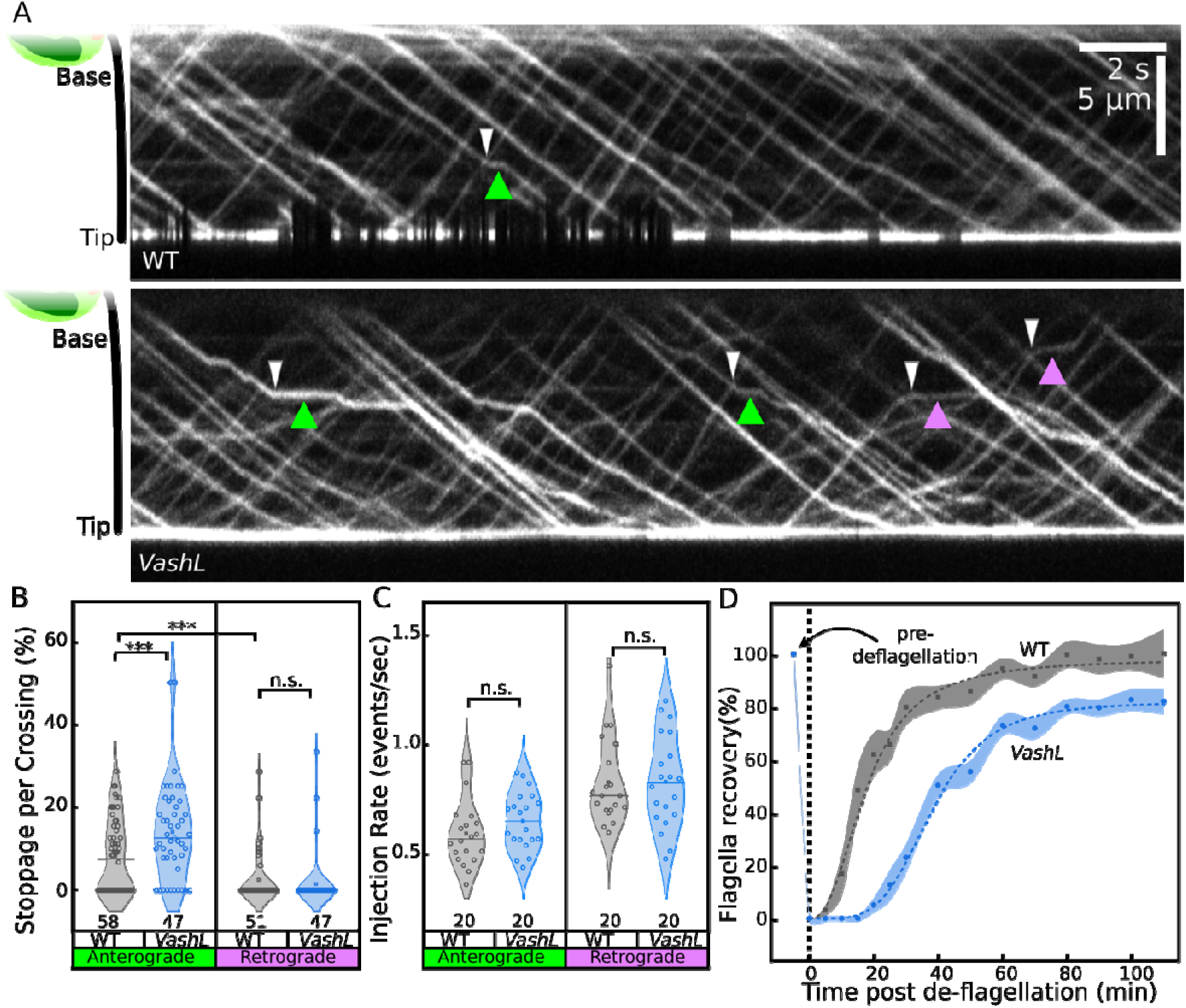
IFT motility in *VashL* cells is frequently interrupted by stoppages. **A)** Kymographs of wild-type (WT, top) or *VashL* (bottom) IFT trains. In *VashL* cells, anterograde or retrograde train stoppages (green or magenta arrowheads respectively) often occur after crossing events (white arrowheads, see **Fig. S3** for additional examples). **B)** Violin plots of stoppages per crossing in wild-type (grey plots) or *VashL* (blue plots) cells. Each datapoint represents an individual train. Horizontal line represents median of each dataset. Pooled data from N = 2-3 independent experiments. Statistics by Wilcoxon Signed rank test. n.s., p-value not significant. ***, p-value < 0.01. **C)** Violin plots of train injection rates from base or tip in wild-type (grey plots) or *VashL* (blue plots) cilia, respectively. Horizontal line represents median of each dataset. Pooled data from N = 2-3 independent experiments. Statistics by Student’s two-tailed t-test. n.s., p-value not significant. **D)** Kinetics of cilia recovery in wild-type (grey plot) or *VashL* (blue plot) cells, expressed as % w.r.t. cilia length pre-deciliation. Error bands represent 95% confidence intervals at each point and dotted lines represent logistic curve fits. N = 12-35 cells for each time point, pooled data from 2 independent replicates for each case.

To further test if the observed alteration of IFT dynamics also leads to perturbations of cilia assembly rate, we measured the rate of cilia regrowth after deciliation in wild-type and *VashL* cells. We found that the initiation of cilia regrowth in *VashL* cells was indeed delayed by about 15 min and progressed at slower rates than in wild-type cells, with mutant cilia never reaching the full length within the experimental time frame (**Fig. 1D**, **Fig. S2C**). Thus, depletion of tubulin detyrosination from axonemal microtubules induces recurrent IFT train stoppages and a reduced growth of cilia.

### Depletion of detyrosinated tubulin does not affect the axonemal structure

To examine the potential impact of detyrosination depletion on the axoneme structure, we utilized cryo-electron tomography and subtomogram averaging to compare wild-type and *VashL* axonemes. The comparison of the 3D structures of the microtubule doublet 96 nm repeat revealed no architectural changes or defects in A-tubules, B-tubules, and major axonemal components (**Fig. S4**). These results show that compromised tubulin detyrosination does not disrupt the assembly of a proper axonemal structure assembly. Thus, the observed cilia growth and IFT motility defects likely stem from a direct effect of the depletion of tubulin modification on the IFT trains and their motors, rather than microtubular or axonemal structural anomalies.

### Reconstitution of IFT train motility *ex vivo* on synthetically polymerized microtubules

To test whether the train stoppage events observed *in vivo* are caused by a direct effect of the tubulin detyrosination depletion, we developed an *ex vivo* approach where we reconstitute the motility of detergent-extracted IFT trains from cilia of *Chlamydomonas* cells on *in vitro* polymerized, polarity-marked microtubules (**Fig. 2**, Materials and Methods). Specifically, *in vitro* polymerized microtubules were applied onto a coverslip and imaged on a TIRF microscope. *Chlamydomonas* cells expressing fluorescently tagged IFT proteins were overlaid onto the microtubules in a droplet of ATP-containing buffer, which supported IFT motility (**Fig. 2A**). A capillary micropipette filled with membrane solubilizing agent (1% Igepal CA-630, henceforth ‘detergent’) was immersed in the droplet and centred over the field of view using a stage-mounted micromanipulator (**Fig. S6A**). A micro-dose of detergent was delivered over the *Chlamydomonas* cells using a calibrated microinjector pump to gently demembranate the cilia and efficiently extract IFT trains (**Fig. S5**; **Fig. S6D, S6E, Video S2**). The detergent shot from the capillary pipette caused rapid demembranation of the cilia, but did not affect the membrane of the cell body, as it is protected by the algae cell wall. Further, as the effective bulk dilution of the detergent was more than 1:10^9^ (20-40 fL (**Fig. S5**) in 5-10 μL of droplet) cilia of cells outside the area of effect were not demembranated. This enabled repeated detergent deliveries in multiple areas within the same droplet, improving throughput and data acquisition. Once extracted from the cilium, IFT trains landed and processively moved along the *in vitro* polymerized microtubules.

**Figure 2.**
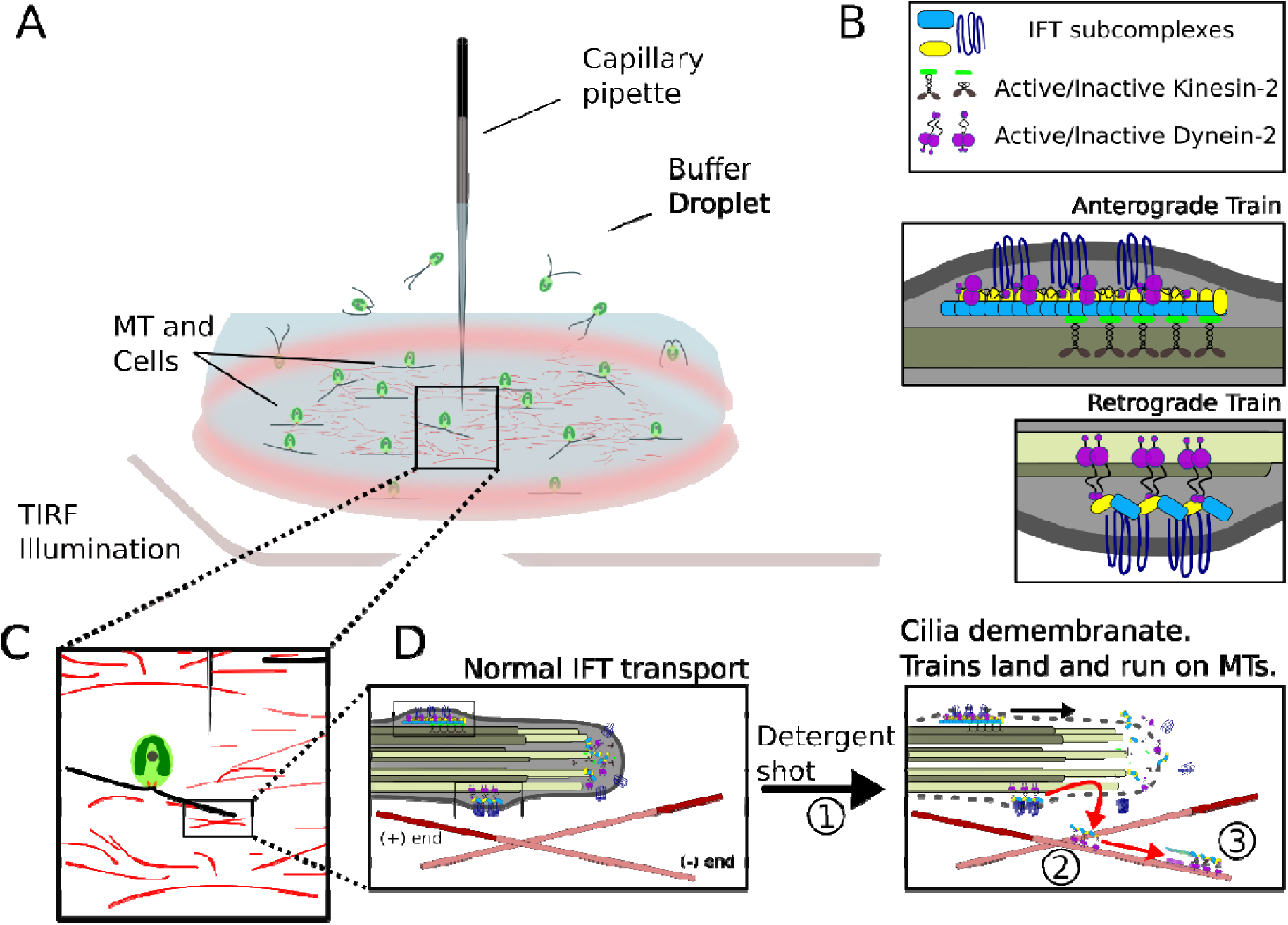
TIRF microscopy coupled with targeted cilia demembranation to reconstitute IFT train motility *ex vivo.* **A)** Illustration of the reconstitution setup. *Chlamydomonas* cells are overlaid on synthetically polymerized microtubules adhered to a coverslip in a droplet of motility buffer on a TIRF microscope. A detergent back-filled capillary micropipette is immersed in the buffer from the top and centred using a three-axis micromanipulator. **B)** Legend showing components of IFT trains (**Top**), and illustrative representations of an anterograde (**Middle**) and retrograde train (**Bottom**). **C)** Illustration of relative positions of microtubules, cells, and capillary pipette. **D)** Illustration of *in vivo* IFT train motility (**Left**) and *ex vivo* IFT train motility on polarity-marked microtubules (**Right**). ⍰ Detergent shot from capillary pipette solubilizes ciliary membrane. ⍰ IFT trains are released into the surrounding environment. Some motile trains continue to move (black arrow) on the demembranated cilia (parent axoneme), while ⍰ others land and move (red arrows) on the polarity-marked microtubules (see also **Video S3**).

Typically, the extracted trains (fluorescently labelled IFT-B, IFT46-mNeonGreen) landed on a microtubule, resumed motility, halted after traveling few micrometres (2.7 ± 2.1 µm, mean ± SD, n = 633 trains) and did not resume motility thereafter (**Figs. 3A, S7, Videos S3, S4**). In rare cases, IFT trains detached from the microtubules instead of halting (**Fig. S7**). We hypothesized that train halting *ex vivo* could be caused by microtubule obstacles, such as other ciliary components interacting with the microtubules, or the loss of motor activity outside the ciliary environment. To test for presence of obstacles, we demembranated cilia of different strains expressing fluorescently labelled non-IFT ciliary components FMG1-B [19], RSP3 [20] or ODA6 [21]. No landing or motility events on microtubules were detectable in these cases (**Videos S5-S7**). This showed that we can specifically reconstitute the motility of IFT trains on microtubules *ex vivo* and that non-IFT components from demembranated cilia did not interfere with motility of *ex vivo* IFT trains in our assay.

**Figure 3.**
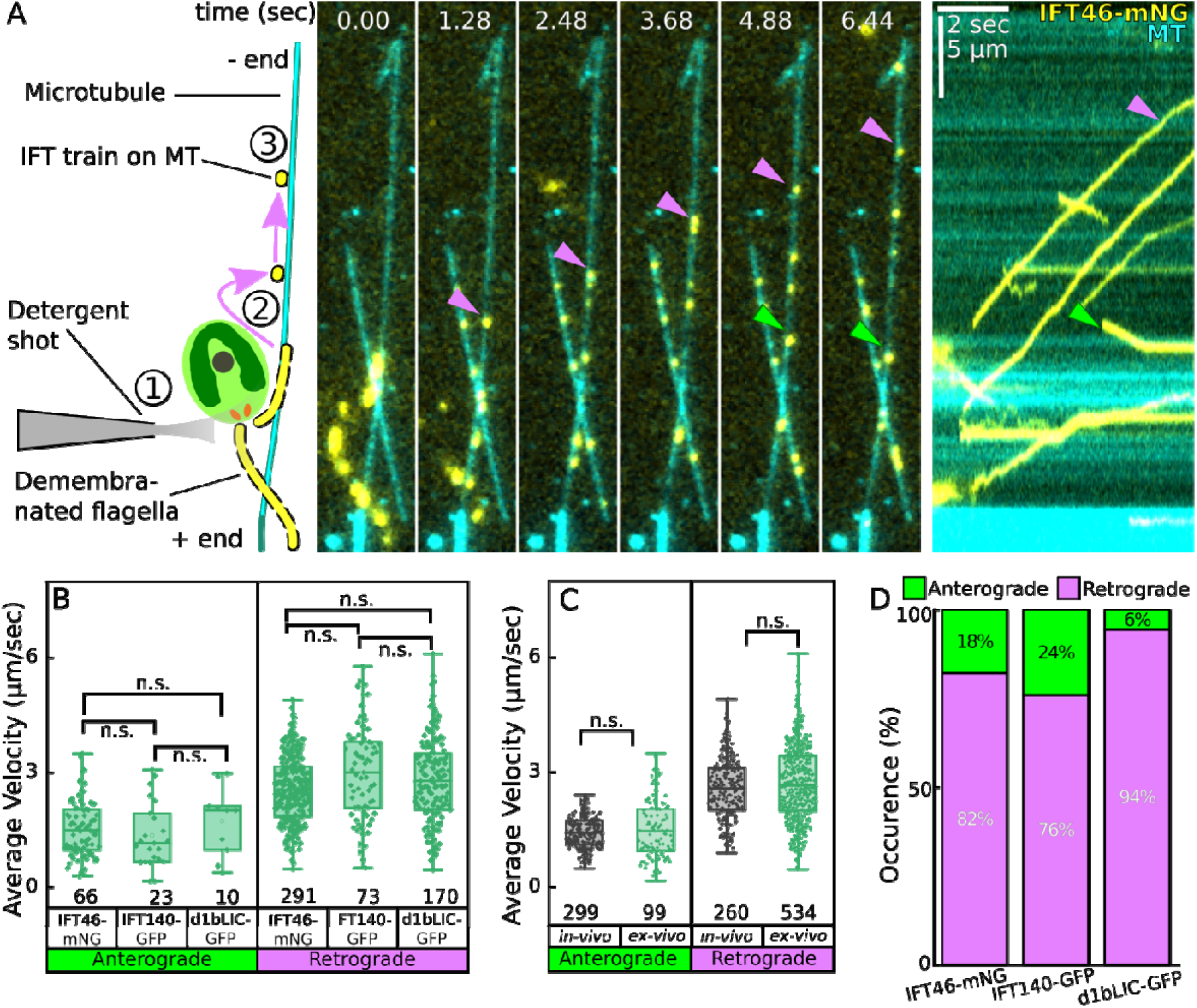
Velocities of IFT trains are conserved during *ex vivo* reconstitution. **A) Left:** Schematic of ⍰ detergent shot, ⍰ landing of IFT train and ⍰ motion on microtubule. **Right:** Representative montage and kymograph of reconstitution of IFT trains (magenta and green arrowheads) comprising mNeonGreen labelled IFT46. The directionality of train movement is determined by plus-end labelling of microtubules (see Materials and Methods, **Video S4**). **B)** Box plots of average velocities as measured from kymographs in (A), for *ex vivo* IFT trains with fluorescently labelled IFT46 (IFT-B), IFT140 (IFT-A), or d1bLIC (IFT Dynein) (see also **Fig. S7**). **C)** Box plots of average *in vivo* (black) or *ex vivo* (green) velocities pooled from (B). For both (B) and (C), statistics by Student’s two-tailed t-test. n.s., p = not significant. **D)** Stacked bar plots of directionality distribution of *ex vivo* trains (labelled with either IFT46-mNeonGreen, IFT140-sfGFP or d1bLIC-GFP).

### All main constitutive components of IFT trains are retained *ex vivo*

To verify whether the extracted IFT trains retained their overall *in vivo* composition, we repeated the reconstitution experiment with strains containing fluorescently labelled IFT-A (IFT140-sfGFP) and IFT-Dynein (d1bLIC-GFP). For all strains, IFT trains were consistently landing and moving on microtubules, demonstrating that all IFT complexes (IFT-A, IFT-B and the motors) are retained in the trains *ex vivo* (**Fig. S8, Videos S8, S9**). The average *ex vivo* velocities of anterograde and retrograde IFT trains were consistent between each strain (**Fig. 3B**) and comparable to the velocities observed *in vivo* (**Fig. 3C**). Additionally, the observed anterograde train velocities were comparable to previously reported single-molecule velocities for the *Chlamydomonas* kinesin-2 complex [22]. Thus, we conclude that *ex vivo* reconstitution could recapitulate *in vivo* IFT train composition and behaviour.

### Anterograde trains have stronger affinity for parent axonemes than retrograde trains and the absence of tubulin detyrosination reduces their affinity for parent axonemes

When quantifying the relative amount of anterograde and retrograde IFT trains on the synthetically polymerized microtubules, we found that a vast majority of *ex vivo* IFT trains moved in retrograde direction (**Fig. 3D**). This suggested that anterograde trains were either less likely to be extracted from cilia or would land less frequently on the microtubules.

To test whether the efficiency of IFT train extraction from cilia was comparable between anterograde and retrograde trains, we demembranated cilia of IFT46-mNeonGreen cells and estimated the number of anterograde or retrograde trains that continued to move along the demembranated cilia (henceforth “parent axoneme”) as fraction of the total number of respective trains in the flagella shaft before demembranation (**Fig. 4, Videos S10, S11**). We observed that anterograde trains were unlikely to detach from wild-type parent axonemes. Only a small fraction of them left the axoneme and landed on the synthetically polymerized microtubules. Contrary to this, retrograde trains had a high detachment probability and often landed on the microtubules (**Fig. 4**). These observations indicate that anterograde trains have a stronger affinity for the parent axoneme than retrograde trains.

**Figure 4.**
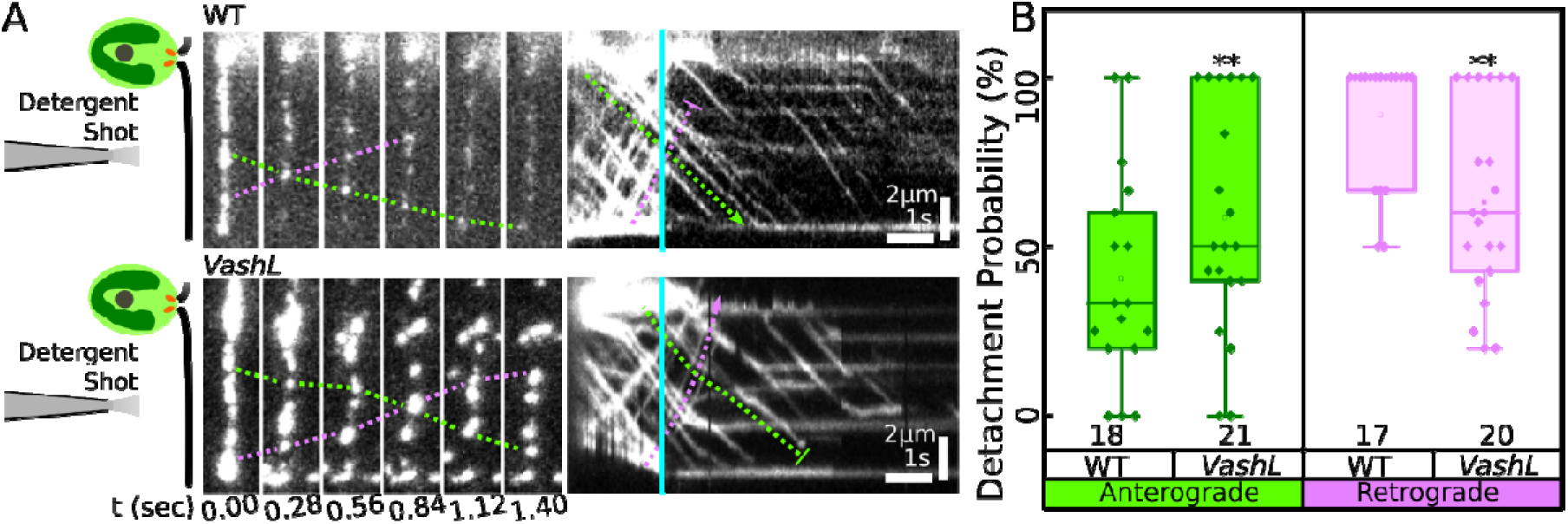
Detachment kinetics of IFT trains from parent axonemes of wild-type or *VashL* cells. **A)** Schematics, representative montage and kymograph of demembranated wild-type (WT) and *VashL* parent axonemes (vertical cyan lines indicate time point of demembranation). In wild-type cells, anterograde IFT trains were less likely to detach from parent axonemes after demembranation (green dashed connectors) as compared to the retrograde IFT trains (magenta dashed connectors). In *VashL* cells, anterograde IFT trains were more likely to detach from the parent axoneme than in wild-type, and retrograde trains are retained more than anterograde trains. **B)** Box plots of detachment probabilities of anterograde (green) and retrograde (magenta) IFT trains in wild-type and *VashL* cells. Statistics by Wilcoxon Signed Rank Test. **, p < 0.05.

Interestingly, when we exposed the detyrosination deficient *VashL* cells to the same detergent treatment, most anterograde trains quickly left the parent axoneme with an increased detachment probability, while the detachment probability was reduced for retrograde trains (**Fig. 4A-B**). In view of these results and because of the recorded high rate of IFT train stoppages in the *VashL* mutant, we propose that depletion of detyrosinated tubulin reduces the affinity of the anterograde IFT trains for the (normally detyrosinated) B-tubules. We suggest that the absence of tubulin detyrosination in *VashL* axonemes interferes negatively with anterograde IFT train affinity and positively with retrograde IFT train affinity.

### Anterograde trains are more likely to land on detyrosinated microtubules and retrograde trains on tyrosinated microtubules

To assess the ability of tyrosinated and detyrosinated microtubules to recruit oppositely directed IFT trains, we estimated the relative train landing probabilities on microtubules with varying amounts of tyrosinated tubulin. To achieve enrichment or depletion of tyrosinated tubulin, brain tubulin was treated with either tubulin tyrosine ligase [23] or carboxypeptidase-A, respectively (Material and Methods). Untreated microtubules were used as controls. To estimate the IFT train landing probability, motile *ex vivo* IFT trains were counted per unit length of microtubule per demembranation event. The landing probability of anterograde trains was found to be higher on microtubules treated by carboxypeptidase-A as compared to untreated microtubules. In contrast, retrograde trains had a higher landing probability on microtubules treated by tyrosine ligase (**Fig. 5A, 5B**). To underscore the preference of anterograde trains for detyrosinated tubulin, and retrograde trains for tyrosinated tubulin, we estimated the relative IFT train landing probabilities with respect to the relative normalized tubulin tyrosination in each case (**Fig. 5C, 5D,** see also **Fig. S9A, S9B**). We find that the landing probability of anterograde IFT trains decreases (**Fig. 5C**), while the landing probability of retrograde IFT train increases (**Fig. 5D**) as function of increasing tubulin tyrosination. Taken together, we suggest that anterograde trains preferentially land on detyrosinated microtubules, while retrograde trains preferentially land on tyrosinated microtubules.

**Figure 5.**
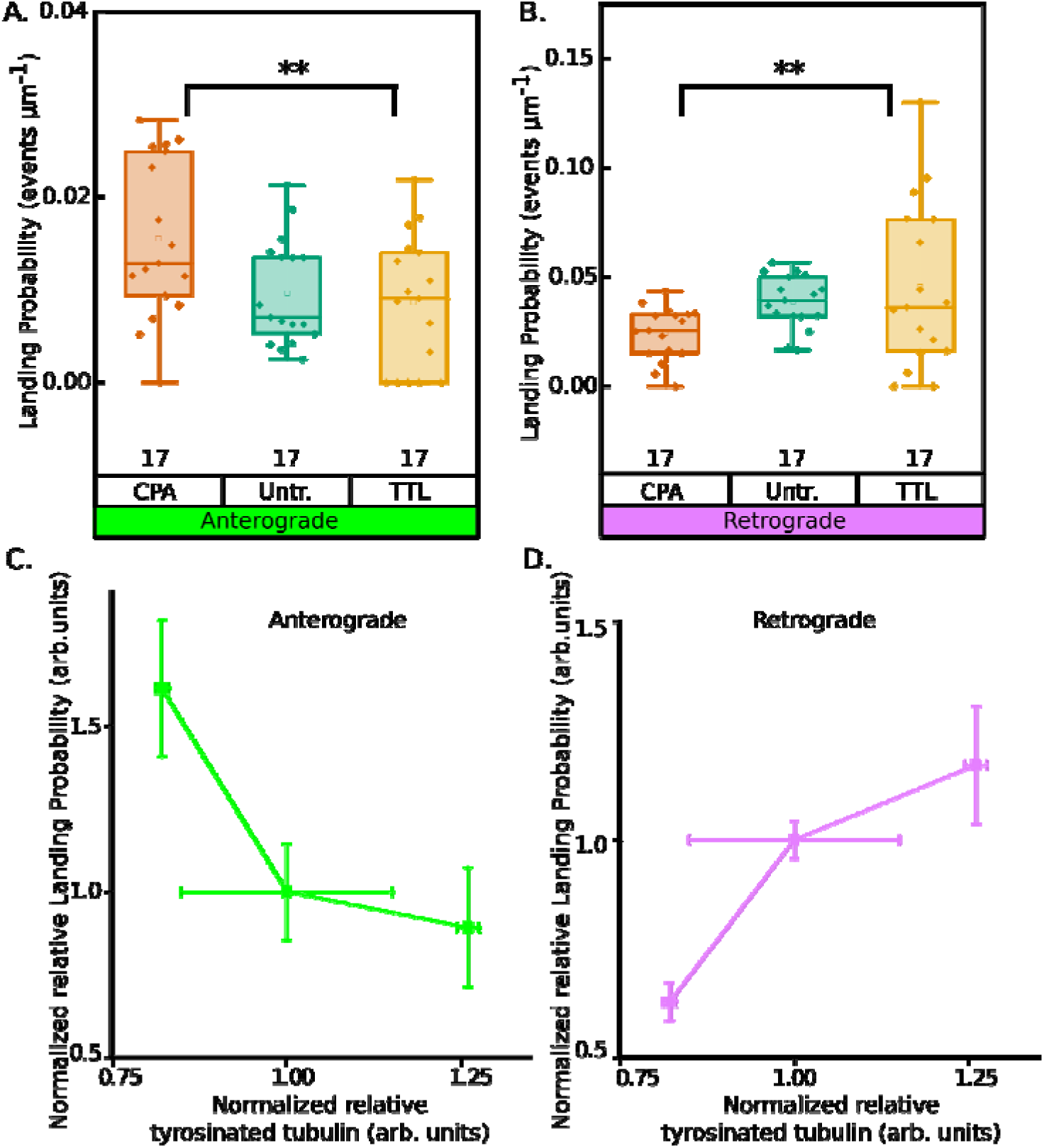
Landing probabilities of *ex vivo* IFT trains on detyrosinated/tyrosinated microtubules. **A)** and **B)** Box plots of landing probability of anterograde or retrograde IFT trains on untreated (green plots), carboxypeptidase-A treated (orange plots), or tubulin tyrosine ligase treated (yellow plots) microtubules. Each dataset is pooled from N = 3-4 independent replicates. Statistics by one-way ANOVA. **, p-value < 0.05. **C)** and **D)** Relative change in anterograde (green plot) or retrograde (magenta plot) IFT train landing probabilities (normalized within samples from **A** and **B**) w.r.t. relative microtubule tyrosination (see **Fig. S9**). N = 3-4 independent replicates for each point. Statistics, one-way ANOVA. **, p-value < 0.05.

### Tubulin Tyrosination/Detyrosination does not affect train velocities

Finally, we measured the velocities of anterograde and retrograde trains on untreated microtubules as well as on microtubules treated with carboxypeptidase-A or tubulin tyrosine ligase. Unlike the landing probabilities, the average train velocities were unaffected on tyrosinated or detyrosinated microtubules (Fig. **S9C**). This further supports the hypothesis that the slowdown of trains observed in *VashL* cells (**Fig. S2B**) is not directly due to a lack of detyrosination, but rather due to stoppages caused by derailment and collisions of oppositely directed trains. (**Fig. 1A, 1B**).

## Discussion

In this work we used CRISPR-mediated targeted mutagenesis to show that depletion of detyrosination in cilia impairs IFT motility and logistics, through temporary stalling of IFT trains when two oppositely directed trains cross on the same microtubule. The recurrent stoppages of anterograde trains led to a discernible slowdown of cilia elongation/regeneration but did not affect the structural integrity of the axoneme. A parallel can be drawn to the phenomena observed in the context of glutamylation and glycylation in *Chlamydomonas* and mouse axonemes, wherein the absence of these specific post-translational modifications does not affect the structure and, despite causing motility phenotypes, does not completely obliterate the beating of the axoneme [6], [7], [8], [24]. We conclude that tubulin post-translational modifications do not orchestrate the process of axonemal structure assembly, but rather the fine regulation of axonemal and IFT constituents.

With the development of our novel method for the reconstitution of the motility of IFT trains *ex vivo*, we demonstrated that tubulin detyrosination and tyrosination have a direct effect on the affinity of anterograde and retrograde trains to the respective microtubules. Anterograde trains showed higher affinity for detyrosinated axonemes in wild-type cells and preferential landing on reconstituted detyrosinated microtubules. Retrograde trains displayed stronger affinity for the tyrosinated axonemes of *VashL* mutant cells and preferential landing on reconstituted tyrosinated microtubules.

Our *in vivo* observations of recurrent stoppage of IFT trains in the *VashL* detyrosination-deficient mutant, leads us to conclude that the differential affinity for detyrosinated and tyrosinated microtubules by IFT kinesin and dynein motors as seen *in vitro* has a specific role in the regulation of IFT logistics in cells. We reason that the mutant phenotype can be attributed to a uniform distribution of tyrosination over both A-tubule abd B-tubule of the microtubule doublets. The consequent loss of specificity for the B-tubule and A-tubule by opposite polarity anterograde and retrograde trains may cause the trains to travel on any microtubule and eventually end up in collisions, leading to the observed stoppage events (**Fig. 6B**). Thus, we propose that in wild-type cilia, despite close physical proximity of both trains to either tubule, asymmetric fidelity of train sorting is maintained by the specific distribution of tyrosinated and detyrosinated tubulin and head-on collisions with oncoming trains are avoided.

**Figure 6.**
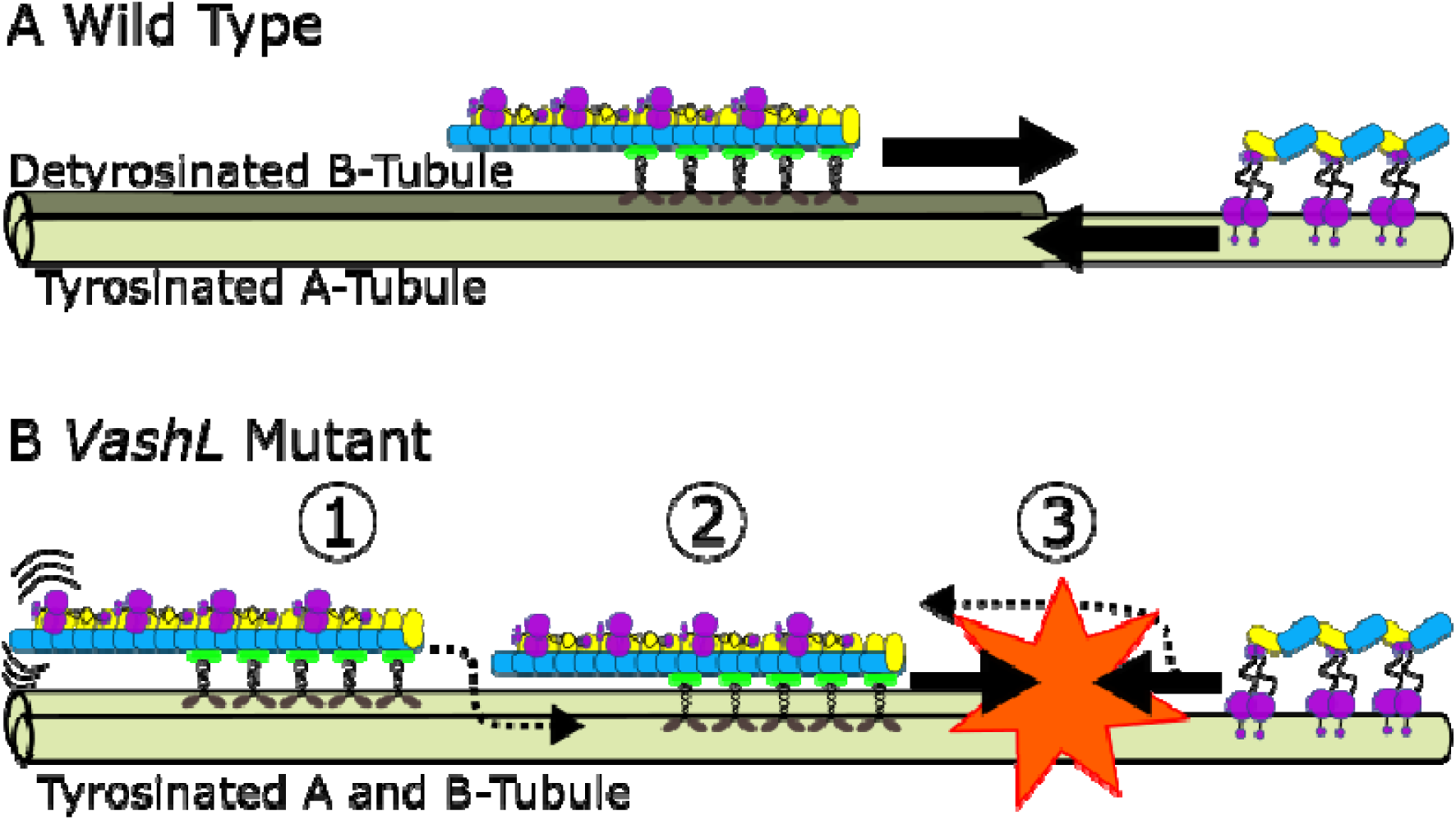
Proposed model for the mechanism of IFT train sorting governed by tubulin tyrosination/detyrosination. **A)** In wild-type cells, anterograde IFT trains get loaded and are retained on the B-tubule (majorly detyrosinated, dark olive). Proximal to the ciliary tip, retrograde IFT trains are conversely loaded and retained on the A-tubule (majorly tyrosinated, light olive), thereby ensuring that train collisions are avoided. **B)** In *VashL* mutant cells, ⍰ anterograde IFT trains do also get loaded onto the B-tubule (tyrosinated, light olive). ⍰ However, due to an increased detachment probability, anterograde IFT trains deviate onto the A-tubule (tyrosinated, light olive). ⍰ Oncoming retrograde IFT trains then collides with missorted anterograde IFT trains leading to stoppage events. An analogous missorting of retrograde trains onto B-tubules is also possible.

Based on our experimental findings, we suggest a novel role of tubulin tyrosination/detyrosination in regulating the dynamics of IFT trains for efficient transport in cilia. We propose a model where train localization is encoded directly on the microtubule tracks. This does not exclude additional regulatory mechanism of microtubule selection by the oppositely directed trains. It has been shown that assembling anterograde trains at the ciliary base associate with the B-tubule already at the transition zone [25]. Thus, in wild-type cells, detyrosination might play a critical role in positively selecting anterograde trains at the ciliary base and then retaining them on the B-tubule along the shaft. At the ciliary tip, instead, the A-tubules extend further than the B-tubules [26]. Thus, retrograde trains might naturally land on A-tubules initially owing to their proximity. We suggest that retrograde trains are then restricted to the A-tubules because they are excluded from the B-tubules due to their lower affinity for detyrosinated tubulin (**Fig. 6A**).

In addition to the direct influence of tubulin post-translational modifications, other mechanisms could facilitate the avoidance of collisions and control tubule-specific transport of anterograde and retrograde IFT trains. For example, single molecule motility assays indicated an adaptive stepping behaviour of ciliary kinesin-2 wherein the intrinsic ability of the motor to switch protofilaments of microtubules was explained by a sidestepping model of IFT train sorting on the microtubule doublets [27]. While direct evidence for sidestepping of rigid linear arrays of multi-motor complexes like a IFT train on microtubule doublets is lacking, the possibility of motor sidestepping functioning cooperatively and redundantly alongside our proposed mechanism for efficient train sorting cannot be ruled out. However, we observed an increased number of stoppage events in *VashL* cells (**Fig. 1B**), where motor sidestepping is likely not supressed. This indicates that sidestepping alone might not be sufficient to avoid the collisions of oppositely directed IFT trains and that tubulin detyrosination is necessary to regulate IFT motility.

Further investigation is required to understand how IFT trains sense the post-translational modifications of axoneme tubulin at a molecular level. Our findings imply that inherent factors within the IFT trains components could read the tyrosination status of the microtubule. Consistent with this idea, *in vitro* studies have shown that mammalian kinesin-1 [6]–[8] and yeast kinesin-2 [15] exhibit increased landing affinity and processivity for detyrosinated microtubules, although such affinity bias has so far not been reported for *Chlamydomonas* IFT kinesin. Additionally, cytoplasmic dynein complexes have shown preferential affinity for tyrosinated microtubules via the CAP-Gly domains of their adaptor dynactin microtubule binding domains [9]–[11]. However, we and others have previously not found any indication of a CAP-Gly domain-like sensor within any component of the IFT complex [2], [28]. Apart from the IFT motors themselves, IFT81/74 [29] and IFT54 [30] have tubulin binding domains. Of these, IFT81/74 is relatively far from the microtubule doublet on anterograde trains [2], and rather binds to soluble Ill/β tubulin dimer as a cargo adapter [29]. It is therefore unlikely unlikely that IFT81/74 reads the detyrosination status of doublet tubules. Also, IFT54 binds directly to both IFT kinesin and dynein [31] and reads only the E-hooks of tubulin or microtubules via its CH-domain [30]. It remains unknown if it senses tubulin tyrosination/detyrosination.

Methodologically we foresee a wide range of applications for our *ex vivo* / *in vitro* approach. While *in vitro* assembly and motility of IFT protein complexes from constitutive proteins has been achieved [29], reconstituting full IFT trains bottoms-up is a formidable challenge. Likewise, although IFT subcomplexes have been isolated using traditional protein isolation methods [32], [33], [34], biochemical purification of entire IFT trains has been unsuccessful so far, mostly due to the complex multimeric nature and distinct structures for anterograde and retrograde IFT trains [33]. IFT trains were seen running only for short periods along the axonemal microtubules of permeabilized primary cilia [35]. Hence, our method, together with the ‘molecular motor toolbox’ [36], presents an opportunity to explore the behavior of native trains in a highly controllable environment to study how other factors, such as viscosity, temperature, and the presence of microtubule-associated proteins [37] and microtubule internal proteins will differentially influence the motility of anterograde and retrograde IFT trains.

## Acknowledgements

The authors would like to acknowledge P. Kiesel and C. Bräuer for technical support, all members of Pigino and Diez labs for fruitful discussions, and the Light Microscopy Facility at MPI-CBG as well as the National Facility for Structural Biology at Human Technopole for technical support. We acknowledge Dennis Diener for comments on the manuscript. Tubulin-Tyrosine Ligase was a kind gift from Michel Steinmetz (Paul Scherrer Institut, Switzerland). The project was supported by the Deutsche Forschungsgemeinschaft (DFG, German Research Foundation) under Germany’s Excellence Strategy – EXC 2068 – 390729961 – Cluster of Excellence, Physics of Life of TU Dresden (starting grant to AC), the European Research Council (ERC) under the European Union’s Horizon 2020 research and innovation program (grant 819826 to GP) and the DFG (grant PI1218/3-1 to GP and SFB 1027 grant to SD). AC was further supported by the Dresden International Graduate School for Biomedicine and Bioengineering (DIGS-BB) and APN by an EMBO long–term fellowship (ALTF number 891-2018) as well as by an HFSP cross-disciplinary fellowship (LT000515/2019).

## Supplementary Figures

**Figure S1.**
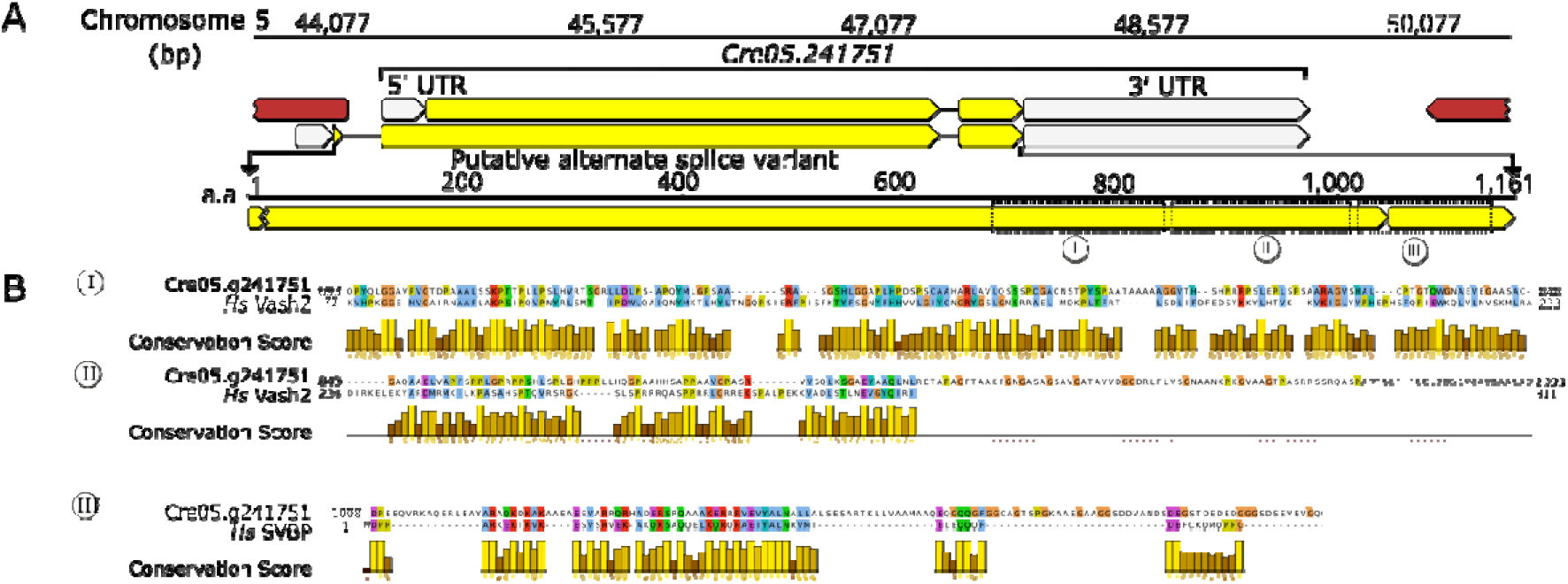
Cre05.g241751 aligns to Human Vash2 and SVBP, related to Figure 1. (**A**) Schematic illustration of *Chlamydomonas reinhardtii* Cre05.g241751 locus on chromosome 5. (**B**) Pairwise sequence alignment of Human Vash2 and SVBP to regions of Cre05.g241751, as marked by arrows.

**Figure S2.**
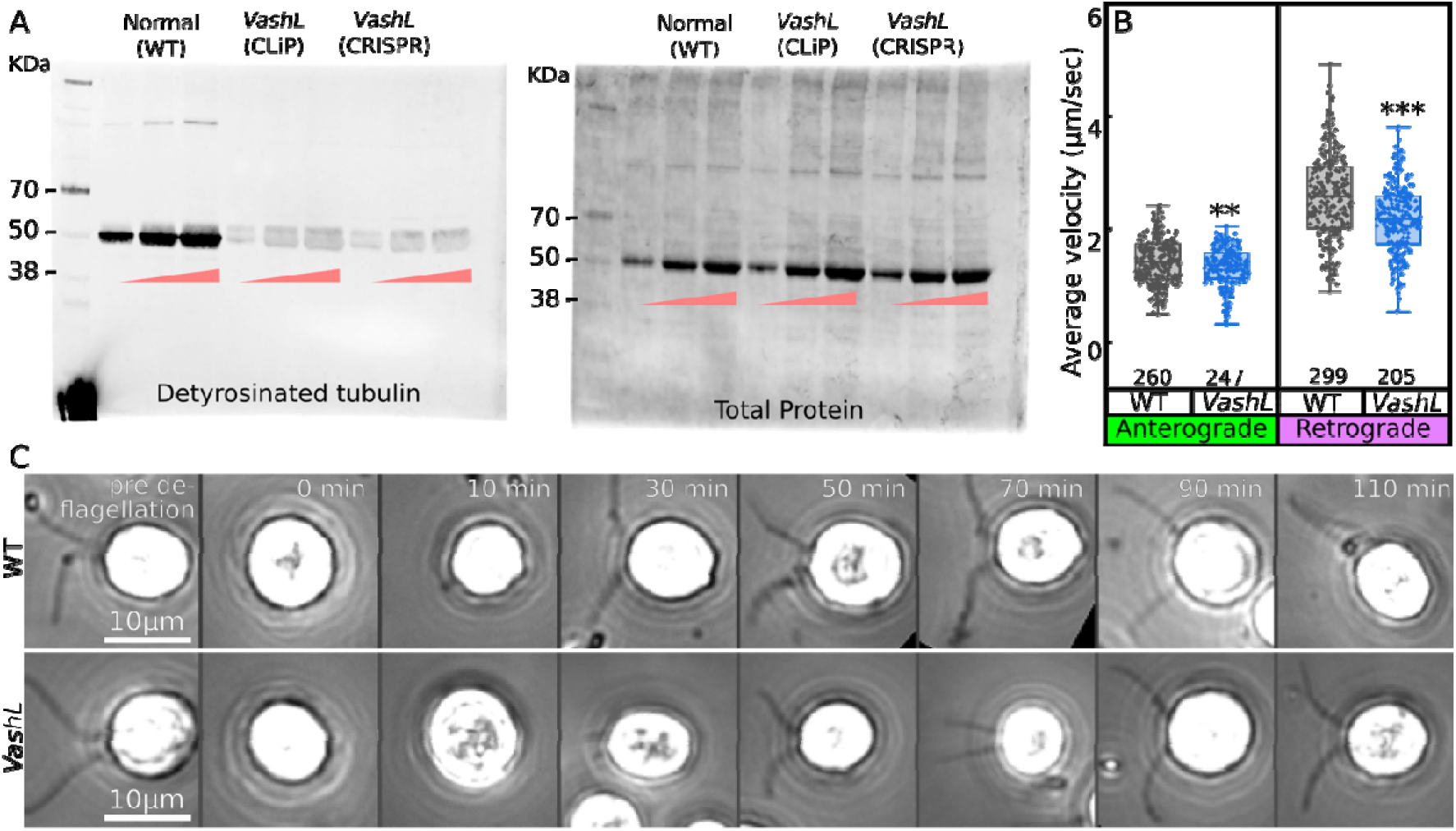
Phenotypic characterization of VashL mutants, related to Figure 1. (**A**) Uncropped blot for detyrosinated tubulin (probed with Merck #MAB5566) (**Left**) and total protein stain (**Right**) of isolated axoneme of either wild-type/IFT46-mNeonGreen), *VashL* CLiP (LMJ.RY0402.233724), or *VashL* CRISPR mutant. Within each group, axonemes are loaded in 1X, 2X or 3X equivalents. (**B**) Box plots of average train velocities in normal (grey) or *VashL* (blue) cells. Pooled data from N = 2-3. Statistics by Student’s two-tailed t-test. **, p-value < 0.05. ***, p-value <0.01. (**C**) Time course snapshots of ciliary recovery after deciliation, in IFT46-mNeonGreen, with (**Bottom**) or without (**Top**) *VashL* mutation. Quantifications in Figure 1D.

**Figure S3.**
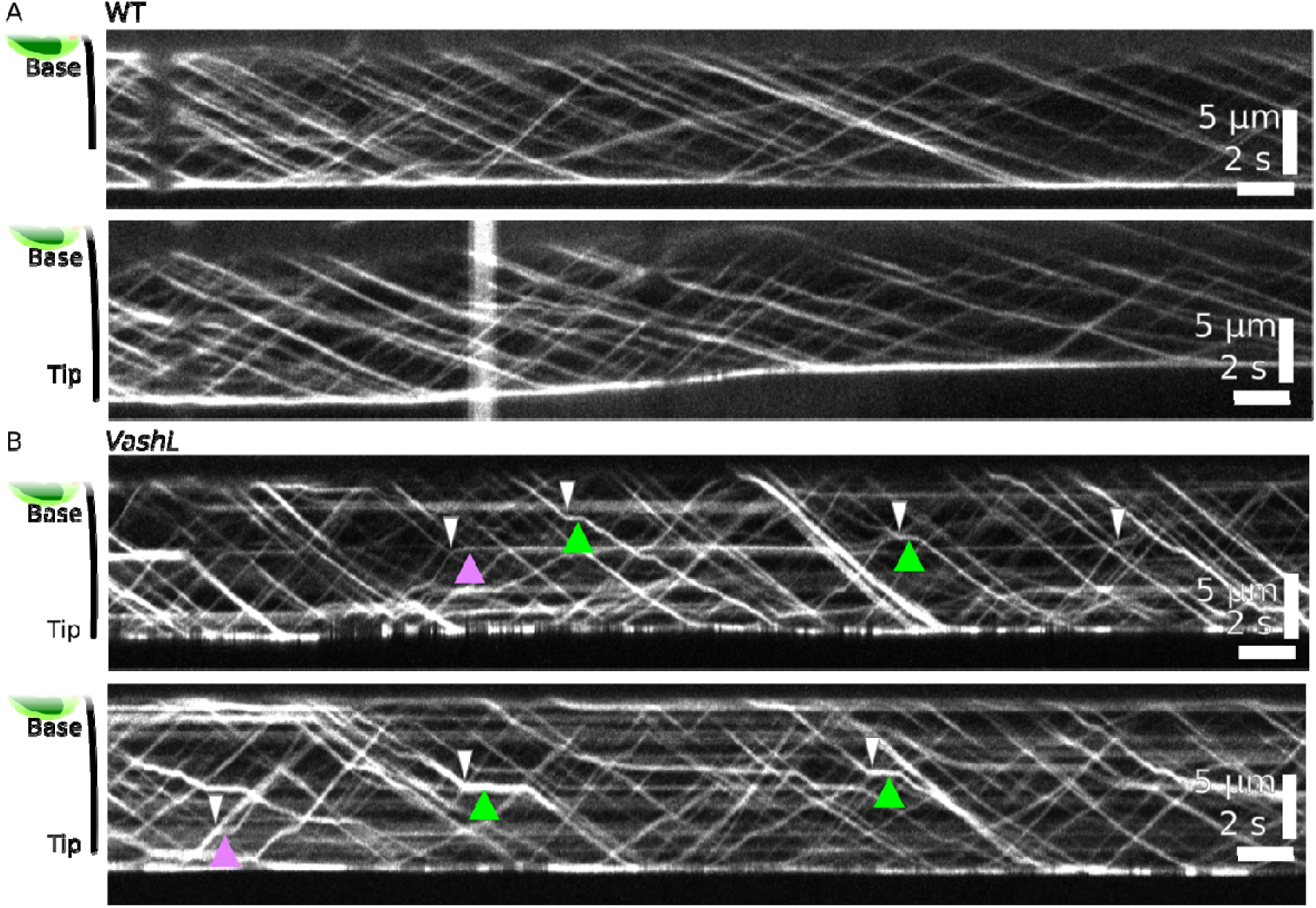
Additional examples of IFT motility in *VashL* cells interrupted by stoppages, related to Figure 1. Kymographs of wild-type (**A**) or *VashL* (**B**) intraflagellar transport (IFT). In *VashL* cells, anterograde or retrograde train stoppages (green or magenta arrowheads) often occur after crossing events (white arrowheads).

**Figure S4:**
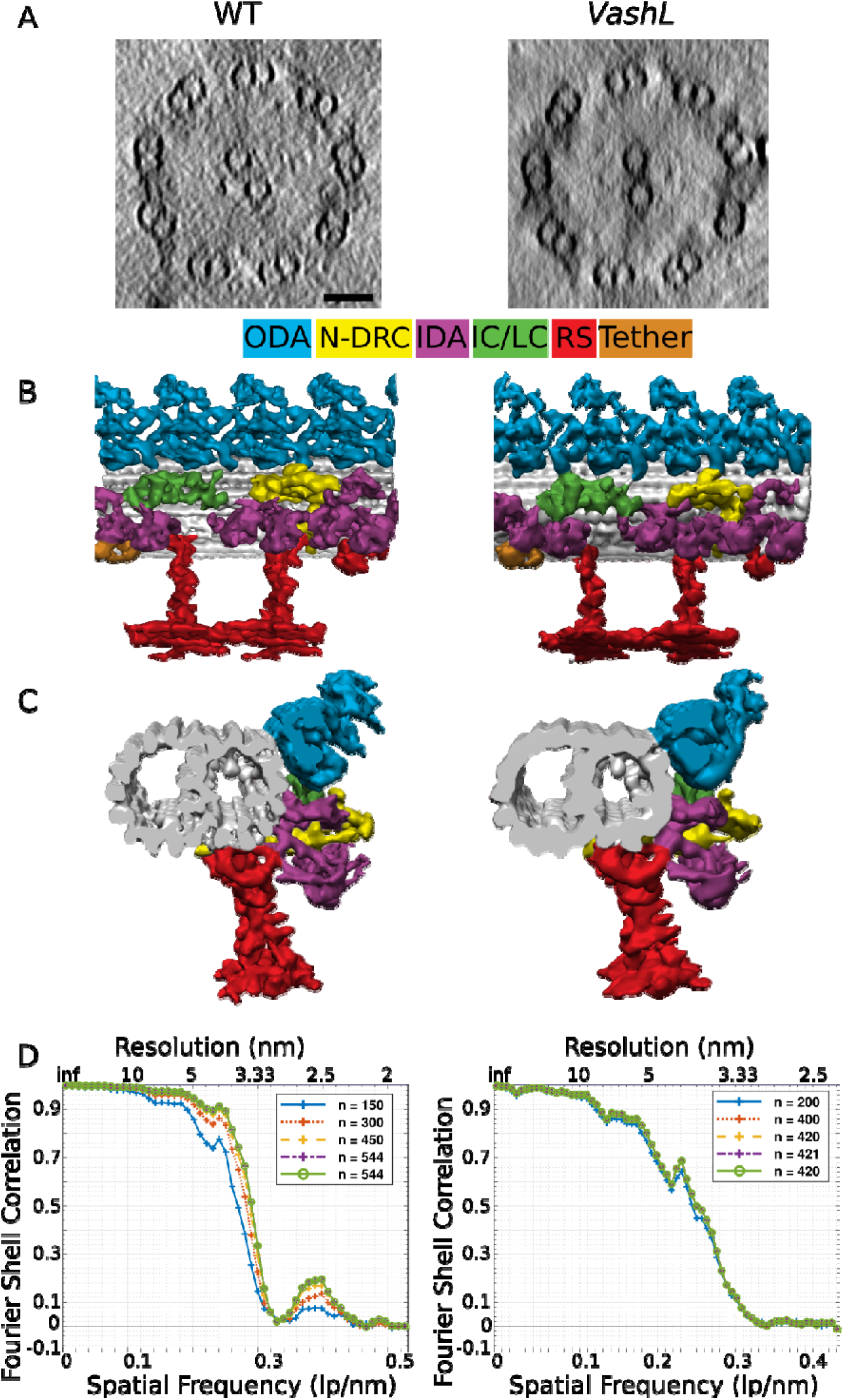
Knock-out of tubulin post-translational modifying enzyme VashL does not alter the macromolecular structure of the Chlamydomonas axoneme. Characterization of the axonemal structure from different Chlamydomonas strains by cryo-electron tomography. (**A**) Tomographic slice (scale bar 50nm), (**B**) and (**C**) Sub-tomogram averaging electron density model of the 96nm-repeat, (**D**) Fourier shell correlation curves for sub-tomogram averaging structures, of a wild-type (**Left**) or *VashL* (**Right**) Chlamydomonas axoneme. Note that, at macromolecular level, the overall structure of the axonemal 96-nm repeat is unaltered in the *VashL* mutant when compared to wild-type.

**Figure S5.**
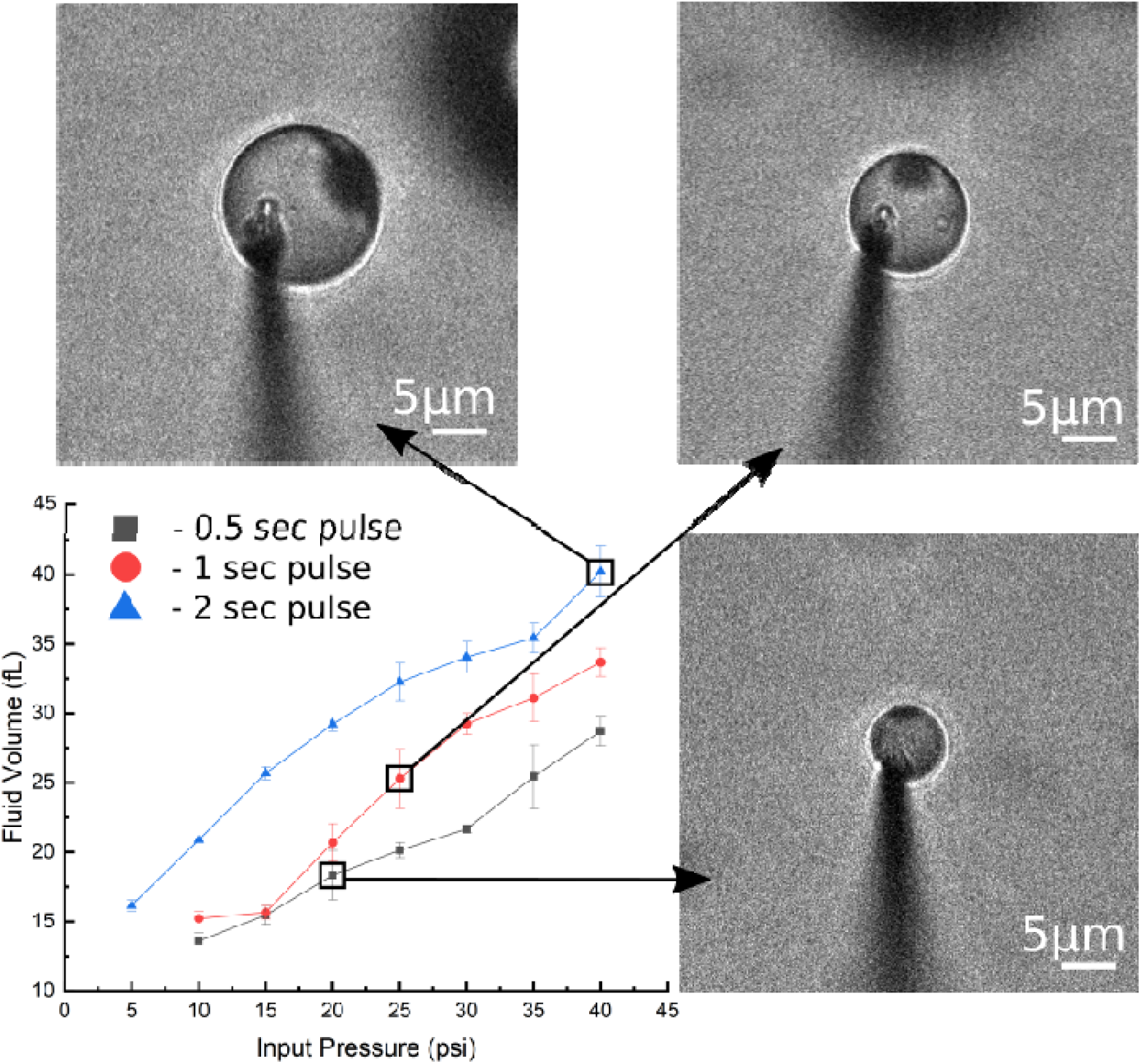
Calibration of microinjector pump. Related to Figure 2 and 3. Volume of ejected fluid (femtoliter) as a function of the input air pressure in injector pump (psi). Volumes estimated from radii of fluid spheres (1.33*π*radius, see Materials and Methods). Stills show representative spheres across the dynamic range of input pressures. Calibrations were performed across N = 3 repeats, average of 3-4 ejections per datapoint.

**Figure S6.**
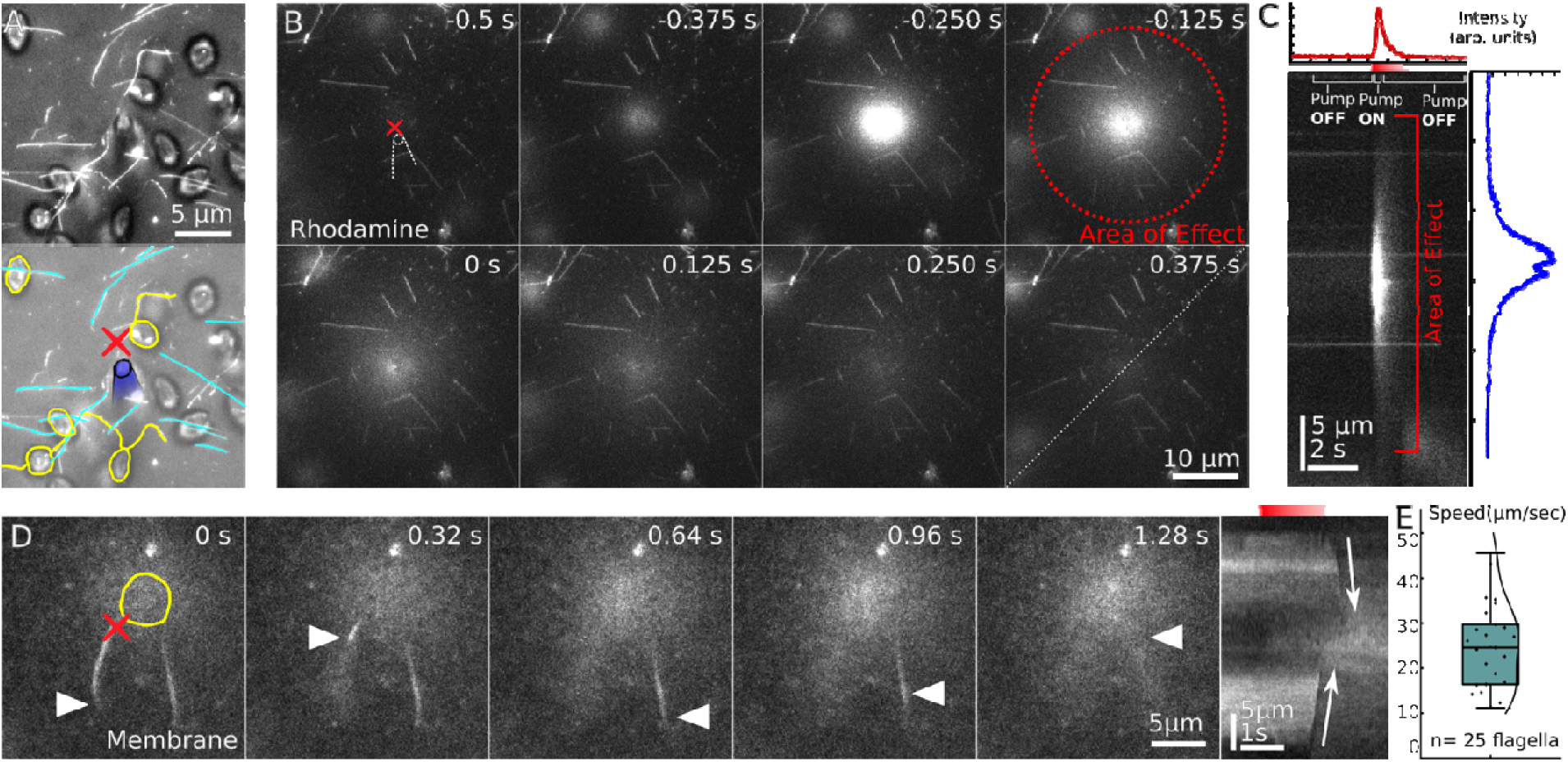
Kinetics of solution ejection and ciliary demembranation, related to Figure 2. (**A**) Representative transilluminated still of a field of view showing relative positions of the *Chlamydomonas* cells (Yellow), microtubules (Cyan), and overhanging capillary micropipette (blue). Red X marks the spot on the coverslip directly under the needle. (**B**) Time series showing diffusion of liquid (dilute Rhodamine) ejected from capillary pipette over time. The force of ejection does not displace microtubules in the neighbourhood. Relative position and centre of capillary pipette as shown. ‘Area of effect’ is a circle of radius 15 μm with X as the centre. See also **Video S1**. (**C**) Kymograph (white diagonal line in (B)) and two-dimension line intensity plots illustrating kinetics of diffusion of ejected liquid. The concentration of the ejected liquid falls exponentially in space and time from the centre of the needle within the area of effect (red square bracket). (**D**) Montage and kymograph of rapid ciliary demembranation (white arrowheads). Cilia proximal to the centre of the capillary (red X) demembranates first followed by the distal cilia. Cell body traced in yellow for reference. See also **Video S2**. (**E**) Single box plot of sample of the speed of demembranation of cilia as shown in (D).

**Figure S7.**
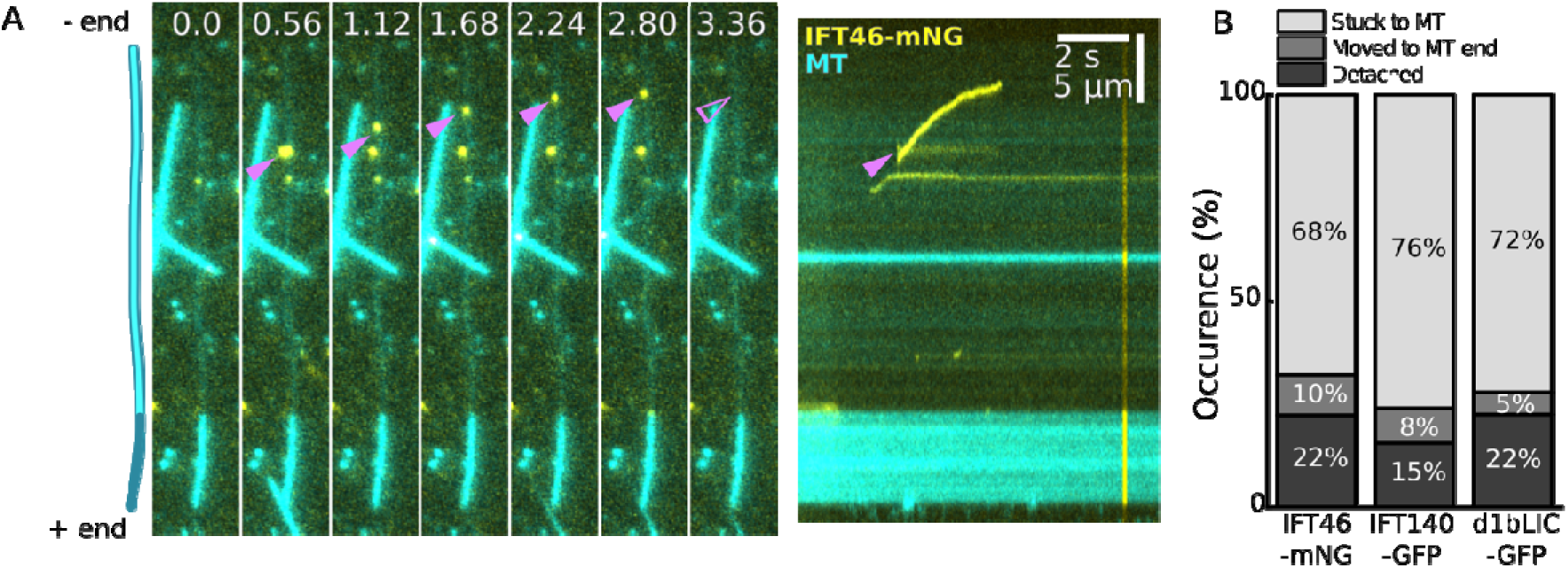
Ex vivo train behaviour on microtubules. Related to Figure 3. (**A**) Montage and example of an ex vivo train that does not halt, but rather detaches from microtubule. (**B**) Percentages of ex vivo train behaviour. Major proportion trains halt on microtubules consistently across multiple samples while a minority detach before halting.

**Figure S8.**
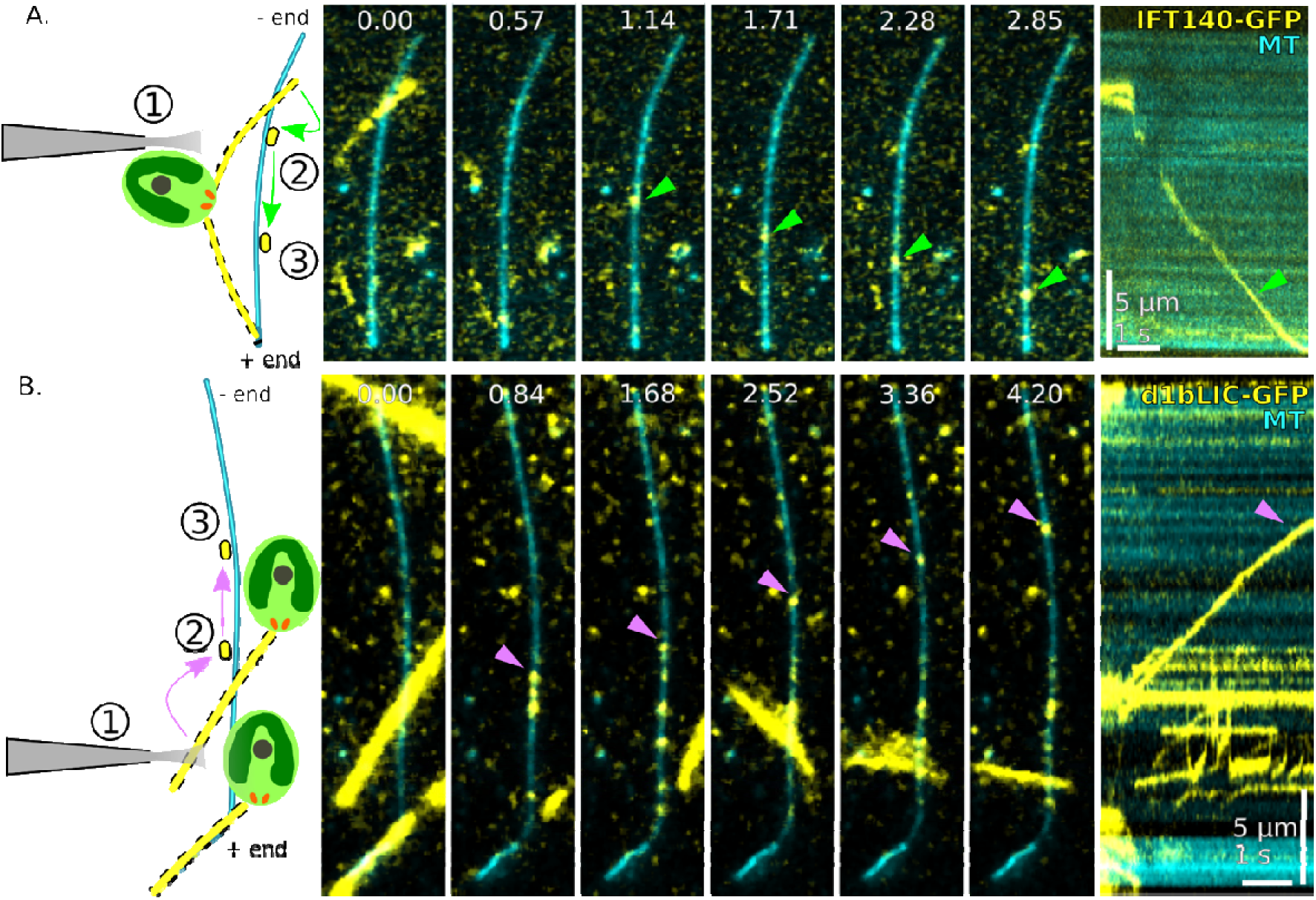
Representative *ex vivo* motility events of IFT trains with labelled IFT-A and IFT-Dynein Complexes. Related to Figure 3. (**A**) **Left:** Schematic of ⍰ detergent shot, ⍰ landing of IFT train and ⍰ motion on MTs. **Right:** Representative montage and kymograph of reconstitution of IFT train (green arrowhead) from IFT140-sfGFP cells. See also **Video S8**. (**B**) **Left:** Schematic of ⍰ detergent shot, ⍰ landing of IFT train and ⍰ motion on MTs. **Right:** Representative montage and kymograph of reconstitution of IFT train (magenta arrowhead) from d1bLIC-GFP cells. See also **Video S9**.

**Figure S9.**
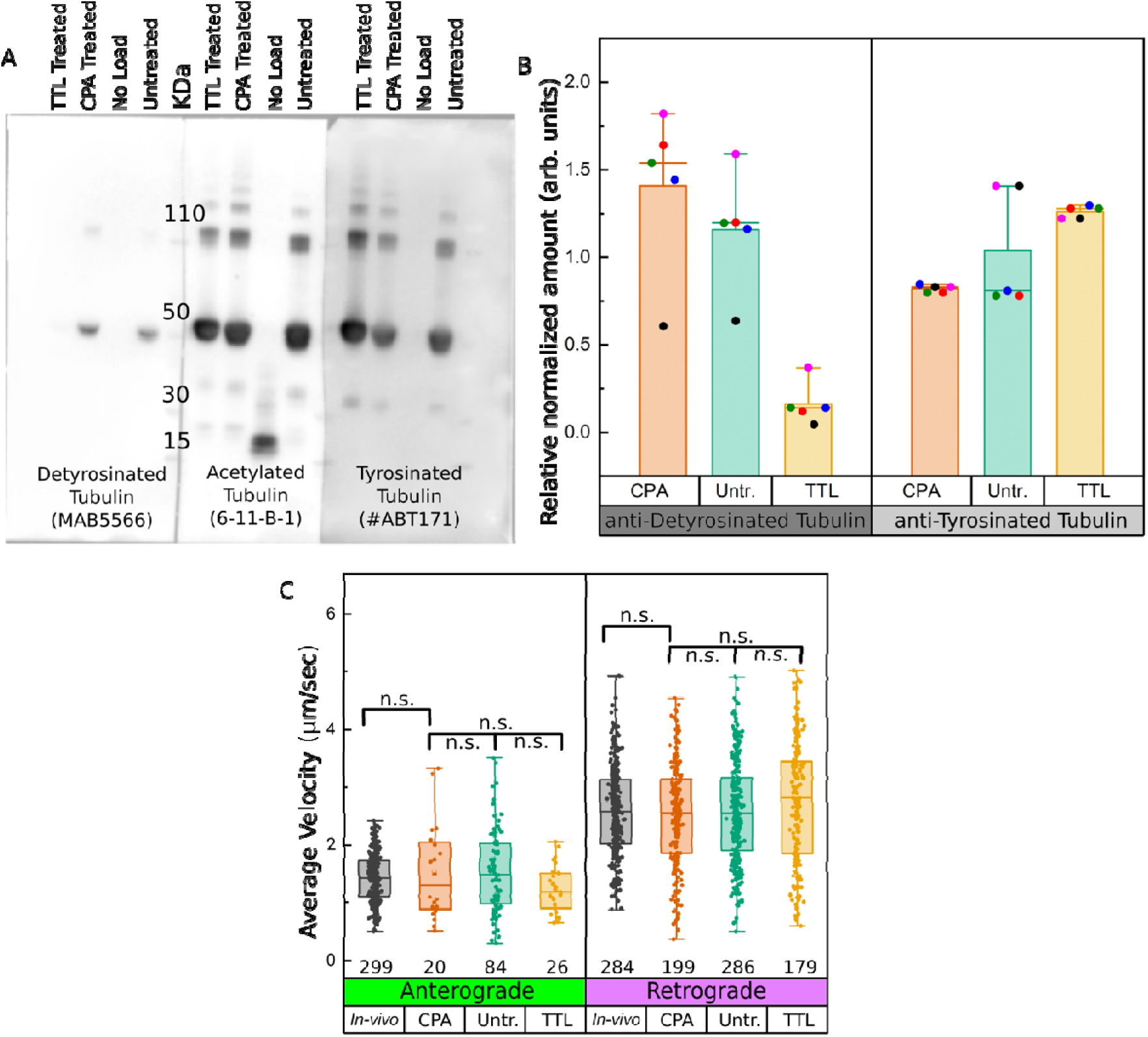
Quantitation of modifications and motility characterization of IFT trains on tyrosinated and detyrosinated microtubules. Related to Figure 5. (**A**) Full uncropped Western blots of identical batches of untreated, carboxypeptidase-A, or tubulin tyrosine ligase treated porcine brain tubulin. Each blot is probed with antibodies specific to either detyrosinated, acetylated or tyrosinated tubulin as mentioned (see Materials and Methods). (**B**) Quantitation and normalization of detyrosinated/tyrosinated tubulin in (A). Samples were normalized w.r.t. acetylated tubulin and further normalized w.r.t. untreated tubulin. Colour scheme in data points of each bar plots represents individual experimental replicates (N=5). (**C**) anterograde or retrograde *ex vivo* average train velocities on microtubules polymerized from treated tubulin obtained in (A). N = 3-4 for all datasets. Statistics by one-way ANOVA. n.s., p-value = not significant

## Supplemental Videos

**Video S1:** Uniform diffusion of ejected liquid (dilute Rhodamine) over the area of effect does not displace microtubules in the neighborhood. Related to Figure 2 and Figure S6.

**Video S2:** Movie of rapid solubilization of Chlamydomonas ciliary membrane caused by detergent shot (indicated by white circle). Related to Figure 2 and Figure S6.

**Video S3:** Representative movie of a experiment showing landing and motility of IFT46-mNeonGreen labelled trains (yellow) on Alexa647-labelled microtubules (cyan; not polarity marked). Cell bodies (cyan; autoflourescence) are shown for clarity. Related to Figure 2 and Figure 3.

Video S4: Representative TIRFM movie used for determining directionality and velocity of ex vivo IFT46-mNeonGreen labelled trains (yellow), moving on TAMRA-labelled microtubule (cyan; bright (+) end). Related to Figure 3.

**Video S5:** Representative TIRFM movie with FMG1b-mNeonGreen labelled (yellow) cells showing no motile events on TAMRA-labelled microtubule (cyan; non polarity marked) upon detergent shot.

**Video S6:** Representative TIRFM movie with pf14::RSP3-NeonGreen labelled (yellow) cells showing no motile events on TAMRA-labelled microtubule (cyan; non polarity marked) upon detergent shot.

**Video S7:** Representative TIRFM movie with oda6::IC2-mNeonGreen labelled (yellow) cells showing no motile events on TAMRA-labelled microtubule (cyan; non polarity marked) upon detergent shot.

**Video S8:** Representative TIRFM movie used for determining directionality and velocity of ex vivo IFT140-sfGFP labelled trains (yellow), moving on TAMRA-labelled microtubule (cyan; bright (+) end). Related to Figure 3 and Figure S8.

**Video S9:** Representative TIRFM movie used for determining directionality and velocity of ex vivo d1bLIC-GFP labelled trains (yellow), moving on TAMRA-labelled microtubule (cyan; bright (+) end). Related to Figure 3 and Figure S8.

**Video S10:** Representative TIRFM movie of IFT46-mNeonGreen labelled cells showing continued association and motility of anterograde train (green arrow) on demembranated parent axoneme, as well as detachment of retrograde train (magenta arrow). Related to Figure 4.

**Video S11:** Representative TIRFM movie of VashL IFT46-mNeonGreen (*VashL* mutant) cells showing detachment of anterograde train (green arrow) from demembranated parent axoneme, as well as continued association and motility of retrograde train (magenta arrow). Related to Figure 4.

## Materials and Methods

### Chlamydomonas Cell Culture

*d1bLIC*::D1bLIC-GFP (CC-4488), *oda6*::IC2-NG (CC-5857), LMJ.RY0402.233724 were purchased from Chlamydomonas Resource Center. FMG1B-mNeonGreen (CC-6012) and *ift46*::NIT IFT46-mNeonGreen (+) (CC-5900) was grown from lab stock. *ift140-1::NIT1 IFT140-sfGFP, pf14::*RSP3-NG were kind gifts from Karl Lechtreck, University of Georgia. Cells were inoculated from TAP-agar plated into liquid TAP medium grown with continuous bubbling of sterile air and normal 12h:12h light-dark cycle.

For de-membranation experiments, ∼5-10 mL cells were harvested, and washed thrice with TAP containing 1 mM EGTA. For final mounting, a small drop (∼2-4 μL) of double-stabilized microtubule suspension was overlaid with cell suspension (final cell density ∼10^6^ /mL), in neutral TAP, with 10 μM paclitaxel and 1 mM ATP final.

### Creation of VashL IFT46-mNeonGreen

*ift46*::NIT IFT46-mNeonGreen (+) is a back-cross of CC-5900 and CC-125. The Chlamydomonas Vash2-SVBP homologue *VashL* (Cre05.g241751_4532) was disrupted by CRISPR/Cas9 insertional mutagenesis with a cassette targeted to the CRISPR guide ATGTGATACCGGCACCACTGTGG on the first exon about one kilobase downstream of the start codon. The cassette is composed of 50 bp homology arms up- and downstream of the cut side flanking a nourseothricin acetyl transferase (NAT) gene under the control of a RbcS2 promoter-terminator pair. Transformation of the *ift46*::NIT IFT46-mNeonGreen (+) was performed as previously described [38]. In brief, cell walls were removed with three incubations in gamete autolysin over 2 hours. During the final round of autolysin incubation, cells were heat shocked at 40°C for 30 minutes, followed by three washes in TAPS medium (TAP+40 mM sucrose). 10^6^ cells in 50uL were mixed with 5uL RNPs at 5uM and 1500uL of BspQI digested and purified donor plasmid. The transformation mixture was electroporated in a 10uL Neon electroporator at 2300V, 12ms, 3 pulses and ejected into 1mL TAPS for overnight recovery. Finally, cells were selected on 1.5% TAP-agar plates supplemented with 7.5ug/mL nourseothricin. Resulting colonies were picked into a 96-well plate filled with TAP and screened for a correct insertion by PCR and sequencing using primers flanking the cut site.

### Coverslip Preparation

22 mm * 22 mm Glass coverslips (Menzel-Glaser, #1.5) were cleaned by sonication in Mucasol/water (1:20; v/v) for 15 min followed by rinsing in deionized water for 2 min. Further, coverslips were soaked in 0.05% solution of Dichlorodimethylsilane (Sigma) for 60 min, rinsed and sonicated twice with 100% methanol, and twice with Milli-Q for 10 min each, and blow dried with nitrogen gas.

### Capillary pipette preparation and manipulation

Capillary pipettes were prepared from thin borosilicate capillaries (BF100-50-15, Sutter Instruments) using micropipette puller (P-1000, Sutter Instruments) as per standard manufacturer protocol. Pipette orifice diameter was maintained at about 2 μm. Taper length of neck was 6-8 mm to ensure sufficient pipette flexibility. Prepared capillary pipettes were back-filled with 1% Igepal CA630 solution (∼5 μL) and manipulated with a combination of coarse (MN-4, Narishige, Japan) and fine (MMO-4, Narishige, Japan) manipulators. Solution ejection parameters (ejection pressure, time pulse) were controlled by a micropipette pump (PV-850, World Precision Instruments, UK).

#### Pipette centring

Vertically mounted capillary pipette was carefully centred and focused to the field of view while imaging live through 10X, 40X, and finally 100X objective using the drive unit of micromanipulator. The working position of the pipette was ∼10-20 μm above the glass surface. Ejection volume calibration was performed according to capillary manufacturer protocols, by ejecting solution into paraffin oil and estimating the volume of the sphere thus obtained. Ejection parameters were ramped as per manufacturer’s recommendation to tune optimal desired ejection volumes. The area of effect for a capillary with orifice diameter of approximately 2 μm was limited to a circle with a radius of roughly 15 μm (Fig. S6B, S6C, see also Video S1).

### Microtubule Polymerization and polarity labelling

Tubulin was purified from porcine brain using established protocols as described previously [39] Polarity-marked microtubules were prepared by preferably extending the plus-ends of long GMPCPP seeds in the presence of N-ethylmaleimide (Sigma) modified tubulin (NEM-tubulin). Briefly, long dim seeds were polymerized from 1:9 rhodamine labelled tubulin (final conc. 2 μM) in the presence of 1 mM GMPCPP (NU-402, Jena Bioscience) in BRB80 at 37°C. For plus end labelling, an extension mix comprising 4 μM 1:3 rhodamine labelled tubulin, 1 μM fresh NEM-tubulin (40 μM unlabelled tubulin incubated with 0.4 mM NEM on ice for 10 min and excess NEM quenched with 20 mM DTT for 10 min), 1 mM GTP(Jena Biosciences) and 4 mM MgCl2 in BRB80 was assembled on ice, warmed at 37 °C for 1 min and incubated with 1/10^th^ volume of dim seeds at 37 °C for 60 min. Microtubules were stabilized with 10 μM taxol in BRB80 (Paclitaxel, Sigma) and harvested by spinning at 17,000 g for 15 min. Polarity-marked microtubules were used on the same day of preparation.

### Enzyme treatment of microtubules

For tyrosination, previously described protocol [23] was adopted with some modifications. 40 μM 1:9 rhodamine labelled tubulin (final conc. 13.33 μM) was incubated with 5 μM Tubulin-Tyrosine Ligase (from Michel Steinmetz, see Acknowledgements), in presence of 0.1 mM sodium salt of L-Tyrosine (Merck), 150 mM KCl, 12.5 mM MgCl_2_, 1 mM DTT and 2.5 mM ATP in BRB80 buffer. The reaction was incubated at 25°C for 3 hours and stopped by adding 1 mM PMSF. For detyrosination, tubulin stocks were prepared as described before [40] with slight modifications. Briefly, 1:9 rhodamine labelled tubulin (final concentration 72 μM) was incubated with Carboxypeptidase-A (Merck) at a relative ratio of tubulin:enzyme (3:1 w/w) at 37°C for 10 min, in presence of 20% (v/v) glycerol, 2 mM GTP. Reaction was stopped by adding 20 mM DTT. Treated tubulin was cycled once in 2 mM GTP at 37°C to remove residual enzyme, flash frozen in liquid nitrogen and finally stored at −80 °C.

### TIRF Microscopy

Imaging was done on a Ti2 Eclipse (Nikon, Japan) equipped with a 100x/1.49NA TIRF objective (MRD01991, Nikon, Japan) with a galvo-driven orbital TIRF consisting of a FRAP/TIRF combination system (iLas2, Gataca Systems, France) coupled to a multi-laser combiner (VS-LMS-MOT100, Visitron Systems, Germany). Multi-color, simultaneous acquisition was done by EMCCD cameras (iXon life, Oxford Instruments, UK) controlled by VisiView software (Visitron Systems, Germany). Data was acquired as continuous stream at 40ms frame time and 1024 * 1024 pixels at a resolution of 0.086 μm px^-1^. Room and sample temperature were maintained at 22°C throughout the experiments.

#### Analysis of TIRFM data

Each detergent ejection and de-membranation event were manually trimmed from a continuous movie stream in Fiji. Kymographs were reconstructed using Spline ROIs (width = 3 px) over microtubules. Velocities were calculated from kymographs using custom written Fiji macros.

Kymographs of Parent Axonemes: Kymographs were traced over all possible locations of axonemes. The resultant kymographs were aligned using StackReg plug-in in Fiji (http://bigwww.epfl.ch/thevenaz/stackreg/). Finally, the registered stacks were Z-projected for maximum intensity to visualize full coverage of axoneme temporally over the de-membranation period.

#### Calculation of Detachment Probability

For each train type on each parent axoneme, ‘Retention fraction’ for anterograde or retrograde trains was calculated as,

R_f_ = N_pa_/N_c_

where,

N_pa_ = Number of trains per demembranated parent axoneme

N_c_ = Number of trains per cilium before de-membranation

Newly injecting trains after de-membranation (particularly anterograde trains) were disregarded in the above evaluation. Further, ‘Detachment Probability’ was calculated as, P_d_ = (1-R_f_) x 100

### Cilia Isolation

Protocol for Isolation of *Chlamydomonas* cilia using pH shock method has been described in detail elsewhere [41]. Briefly, a bubbling culture of cells was washed in HMDEK buffer (30 mM HEPES, pH 7.4, 5 mM MgSO_4_, 1 mM DTT, 0.5 mM EGTA, 25 mM KCl) thrice. Deciliation was done by adding 0.5 N acetic acid dropwise till pH of the buffer was reduced to 4.5 for 45 to 60 secs, before neutralization to pH 7.0 with 0.5 M KOH. Isolated cilia were suspended in HMDEK buffer containing 1 mM DTT, 1 μM Aprotinin and then were separated from cell bodies by spinning over a sucrose cushion (25% v/v in HMDEK buffer). Axonemes were obtained by treating isolated cilia with 1% Igepal CA-630 solution to solubilize ‘Membrane + Matrix’ fraction.

### Western Blotting

Axonemes isolated from *ift46*::NIT IFT46-mNeonGreen, LMJ.RY0402.233724, and IFT46 mNeonGreen:*VashL* were boiled in 1X Laemmli Buffer and run in equivalents on 4-15% continuous gradient SDS-PAGE gels (Invitrogen). Bands were visualized by Coomassie staining and normalized between samples. To estimate Detyrosinated tubulin, normalized axoneme equivalents were re-run on 4-15% SDS-PAGE gels, blotted onto Nitrocellulose membrane, and probed with anti-Detyrosinated Tubulin antibody (MAB5566, Merck). To estimate relative enrichment of tyrosinated or detyrosinated tubulin after enzyme treatment, 3 identical treated tubulin batches were run on 4-15% SDS-PAGE gels, blotted onto Nitrocellulose membrane. Blots were probed with with either anti-Tyrosinated (ABT171, Merck), Detyrosinated (MAB5566, Merck), or Acetylated Tubulin antibody (6-11B-1, Santa Cruz), developed using chemiluminescence imager (Azure Biosystems).

### Plunge freezing

A Leica Automatic Plunge Freezer EM GP machine was used to perform plunge freezing as it counts with an integrated humidity control chamber. Cu supported 3.5/1 Holey Carbon grids (Quantifoil) were glow discharged on both sides for six seconds each using atmospheric air to render them more hydrophilic. In the meantime, the plunge freezer chamber temperature was set to 32 °C to obtain and maintain high humidity (above 80%). The plunge freezer was set to detect the wetting of the filter paper (Whatman) upon blotting (around 2 seconds blotting time, no automatic plunging), which was mounted on one side only with the purpose of blotting from the grids back side. The sample was loaded on both sides (3.5 μL total) and incubated for 30 seconds before the addition of 1.5 μL of sonicated gold fiducials suspended in PBS, followed by immediate blotting. In many cases automatic blotting was not optimal, requiring swift manual blotting before plunging. Grids were stored in grid-boxes which were themselves stored in cryogenic tanks.

### Cryo-Electron Tomography

Acquisition of cryo-EM images for wild-type axonemes was done with a 300 kV Thermo Fisher Titan Halo TEM with a field emission gun (FEG) electron source and a Gatan K2 Summit direct electron detector with energy filter using a slit width of 20 eV. Acquisition of cryo-EM images for VashL axonemes was done with a 300 kV Titan Krios and a Falcon4i camera with a SelectrixX energy filter with a slit width of 20 eV. To facilitate the location of regions of interest containing well preserved cilia, SerialEM [42] was used to generate a full grid overview by automatically acquiring and stitching low magnification (210x) images. Tilt series were acquired with SerialEM on areas of interest at 30,000x nominal image magnification (33kX for *VashL* dataset), resulting in a calibrated pixel size of 4.72 Å (counted mode) (3.76 Å for the *VashL* dataset). Tilt series were recorded with 2° increments with a bidirectional tilt scheme from –24° to 42° and from −26° to –42° (3° tilt increment and dose symmetric scheme from 0 to ± 60° for the *VashL* dataset. The defocus target varied from – 1.5 to −5.5μm and the cumulative dose was 95- 120 e - per Å 2 per tomogram. Images were acquired in the dose fractionation mode with 0.25 s frame time to a total of around 8 frames per tilt image (.eer format for the *VashL* dataset). Drift was kept below 1 nm/s. The frames were aligned using K2Align software (motioncorr2 for the *VashL* dataset), which is based on the MotionCorr algorithm [43]. Tomogram reconstruction was performed using Etomo from IMOD v.4.10.11-α using weighted backprojection [44]. Contrast transfer function curves were estimated with CTFPLOTTER and corrected by phase-flipping with the software CTFPHASEFLIP, both implemented in IMOD [45]. Dose weighted filtering was also performed using the mtffilter command from the IMOD package. In some cases, a nonlinear anisotropic diffusion filter by IMOD [44] was applied on low-contrast tomograms acquired closer to focus. The *VashL* dataset tomograms were filtered with a deconvolution filter.

### Sub-tomogram averaging

IMOD v.4.10.11-α and PEET v1.11.0 were used for sub-tomogram averaging [46]. Using 3dmod, included in the IMOD package, tomograms were inspected looking for signs of adequate structural preservation, namely, roundness and straightness of the axonemes and presence of unaltered axonemal components such as radial spokes and dynein arms. To model the location of each 96-nm repeat within tomograms, the coordinates of points of the central axis of A-tubules at the base of every RS2 were determined manually and saved in .mod format (supported by IMOD software). At least a thousand of these points were determined per experimental condition. Using PEET (v 1.11.0), each sub-tomogram containing a 96-nm repeat was aligned to a common reference so that all microtubule axes remained parallel to each other with the right polarity. The particles from each microtubule doublet were averaged independently to determine the angles of rotation that should be applied to each microtubule doublet to pre-align it with the common reference, providing a good starting point for automated fine alignment. Missing wedge compensation was used, one single particle was commonly used as an initial reference per experiment. Soft masks outlining the 96-nm repeat structure were used for alignment optimization. The maximum search ranges for translations and rotations of each particle during alignment to iteratively refined references were estimated based on the precision achieved during manual modeling and pre-alignment, normally resulting in values of ±12° around the Y axis, ±6° for the remaining axes and 8 pixels for translations in each direction. This strategy, laborious compared to fully automated sub-tomogram averaging routines, provides human-proofread averages without missing-wedge, chirality, or polarity artifacts. Routinely, the final average was generated discarding the worst 10% of particles based on their cross-correlation coefficient score against the final reference (consisting of the average of the 66% best aligned particles). Visualization of averaged electron density maps was performed in 3dmod from IMOD. 3D rendering of iso-surfaces was performed using UCSF Chimera (v 1.14) [43], [47]

